# Mechanisms for Electron Uptake by *Methanosarcina acetivorans* During Direct Interspecies Electron Transfer

**DOI:** 10.1101/2021.06.11.448114

**Authors:** Dawn E. Holmes, Jinjie Zhou, Toshiyuki Ueki, Trevor Woodard, Derek R. Lovley

**Author notes:** Corresponding author Dawn Holmes. Both authors contributed equally (Dawn E. Holmes, Jinjie Zhou).

## Abstract

Direct interspecies electron transfer (DIET) between bacteria and methanogenic archaea appears to be an important syntrophy in both natural and engineered methanogenic environments. However, the electrical connections on the outer surface of methanogens and the subsequent processing of electrons for carbon dioxide reduction to methane are poorly understood. Here we report that the genetically tractable methanogen *Methanosarcina acetivorans* can grow via DIET in co-culture with *Geobacter metallireducens* serving as the electron-donating partner. Comparison of gene expression patterns in *M. acetivorans* grown in co-culture versus pure culture growth on acetate revealed that transcripts for the outer-surface, multi-heme, *c-*type cytochrome MmcA were higher during DIET-based growth. Deletion of *mmcA* inhibited DIET. The high aromatic amino acid content of *M. acetivorans* archaellins suggests that they might assemble into electrically conductive archaella. A mutant that could not express archaella was deficient in DIET. However, this mutant grew in DIET-based co-culture as well as the archaella-expressing parental strain in the presence of granular activated carbon, which was previously shown to serve as a substitute for electrically conductive pili as a conduit for long-range interspecies electron transfer in other DIET-based co-cultures. Transcriptomic data suggesting that the membrane-bound Rnf, Fpo, and HdrED complexes also play a role in DIET were incorporated into a charge-balanced model illustrating how electrons entering the cell through MmcA can yield energy to support growth from carbon dioxide reduction. The results are the first genetics-based functional demonstration of likely outer-surface electrical contacts for DIET in a methanogen.

**Importance:** The conversion of organic matter to methane plays an important role in the global carbon cycle and is an effective strategy for converting wastes to a useful biofuel. The reduction of carbon dioxide to methane accounts for approximately a third of the methane produced in anaerobic soils and sediments as well as waste digesters. Potential electron donors for carbon dioxide reduction are H_2_ or electrons derived from direct interspecies electron transfer (DIET) between bacteria and methanogens. Elucidating the relative importance of these electron donors has been difficult due to a lack of information on the electrical connects on the outer surface of methanogens and how they process the electrons received from DIET. Transcriptomic patterns and gene deletion phenotypes reported here provide insight into how a group of *Methanosarcina* that play an important role in methane production in soils and sediments participate in DIET.

## Introduction

The pathways for carbon and electron flux in methanogenic environments are of interest because of the biogeochemical significance of methane production in diverse soils and sediments as well as the importance of anaerobic digestion as a bioenergy strategy (1, 2). Diverse communities of bacteria convert complex organic matter primarily to acetate and carbon dioxide that are then converted by methanogenic archaea to methane. Low-potential electrons derived from the oxidation of organic compounds to acetate and carbon dioxide are delivered from the bacterial community to methanogens to provide the necessary reducing power for the reduction of carbon dioxide to methane.

Two fundamentally different mechanisms for this interspecies electron transfer are known. In direct interspecies electron transfer (DIET), electron-donating microbes and methanogens establish direct electrical connections that enable electron transfer from the electron-donating partner to the methanogen to support carbon dioxide reduction (2-4). In interspecies H_2_ transfer, the electron-donating partner transfers electrons to protons, generating H_2_, which functions as a diffusive electron carrier to H_2_-utilizing methanogens, which oxidize the H_2_ to harvest electrons for carbon dioxide reduction (5-7). Formate can also serve as a substitute for H_2_ (6, 8, 9).

The relative importance of DIET and interspecies H_2_/formate transfer in methanogenic soils/sediments or most anaerobic digesters is unknown. Measurements of H_2_ turnover rates in methanogenic soils, sediments, and anaerobic digesters accounted for less than 10% of the electron flux required for the observed rates of carbon dioxide reduction to methane (10-12), suggesting that H_2_ exchange was not the primary route for interspecies electron transfer (13). Those results do not rule out interspecies formate exchange, but rapid exchange between formate and H_2_/carbon dioxide in methanogenic environments prevents accurate assessment of formate fluxes (14). The relatively low reported rates of H_2_ turnover are consistent with DIET, but a method for directly measuring the electron fluxes between cells in complex environments has not yet been developed.

An alternative strategy for elucidating the significance of interspecies H_2_/formate transfer and DIET might be to extrapolate from the composition of the microbial community and transcriptional or proteomic data (15-17). For example, *in situ* gene expression patterns of *Geobacter* and *Methanothrix* species, which were abundant in methanogenic rice paddy soils, suggested that they were participating in DIET (17). The likely participation of paddy-soil *Geobacter* species in DIET could be surmised from high levels of expression for electrically conductive pili (e-pili) and a *c*-type cytochrome known to be important for DIET. High expression of genes for carbon dioxide reduction in *Methanothrix* species, which are unable to use H_2_ or formate as electron donors, indicated that *Methanothrix* species were one of the electron-accepting partners for DIET. However, such analyses are far from comprehensive, in part because the full diversity of microbes that can participate in DIET is poorly understood. New genera of bacteria and methanogens capable of DIET are increasingly being identified (18-20). Furthermore, gene expression patterns diagnostic for ongoing DIET need to be elucidated for microorganisms, such as *Syntrophus* (18) and some *Methanosarcina* species (16, 19, 21-23) that have the physiological flexibility to participate in either DIET or interspecies H_2_/formate transfer.

Comparative transcriptomic analysis of *M. barkeri* growing via DIET versus interspecies H_2_ transfer, revealed potential routes for intracellular electron flux for DIET that employ protein complexes and electron carriers that are also important for the conversion of carbon dioxide to methane with H_2_ as the electron donor (21). Outer-surface electrical contacts for DIET were not definitively identified. *M. barkeri* lacks multi-heme outer-surface *c*-type cytochromes (24) that are important electrical contacts for extracellular electron exchange in some bacteria and archaea (13). *M. mazei*, which like *M. barkeri*, can reduce carbon dioxide with electrons derived from H_2_ or DIET, has a gene for a five-heme, *c*-type cytochrome, but deletion of the cytochrome gene did not negatively impact DIET (19).

*M. barkeri* and *M. mazei* are physiologically classified as Type I *Methanosarcina* (22). Key physiological characteristics of Type I *Methanosarcina* are the ability to use H_2_ as an electron donor for carbon dioxide reduction as well as for energy conservation from the conversion of acetate to methane via intracellular H_2_ cycling. Although Type I *Methanosarcina* can serve as the electron-accepting partner for DIET, they are typically most abundant in high energy environments with relatively fast rates of organic carbon turnover in which H_2_ is more likely to be an intermediate in interspecies electron transfer (22).

In contrast, Type II *Methanosarcina* predominate in more stable, steady-state environments with slower rates of organic matter metabolism likely to favor DIET (22). Key physiological characteristics of Type II *Methanosarcina* include the inability to use H_2_ as an electron donor, energy conservation during acetate metabolism via an Rnf complex, and the presence of an outer-surface multi-heme *c*-type cytochrome that is important for electron transfer to extracellular electron acceptors (22). The inability of Type II *Methanosarcina* to utilize H_2_ or formate as an electron donor for carbon dioxide reduction, but to participate in DIET (22) is expected to simplify the study of their routes for electron flux during DIET.

In order to better understand DIET mechanisms in Type II *Methanosarcina*, we investigated DIET in *M. acetivorans. M. acetivorans* is the most well-studied Type II *Methanosarcina* and is genetically tractable (25-28). Transcriptomic and gene deletion studies (29) demonstrated that its multi-heme outer-surface *c*-type cytochrome MmcA is important for extracellular electron transfer to the humic substances analogue anthraquinone-2,6-disulfonate (AQDS). Here we report that *M. acetivorans* can function as the electron-accepting partner for DIET and provide insights into mechanisms for electron uptake and energy conservation during DIET-based growth.

## Results and Discussion

### *Methanosarcina acetivorans* can participate in DIET

Co-cultures of *M. acetivorans* and *G. metallireducens* metabolized ethanol to methane. As previously observed with co-cultures of *G. metallireducens* and other electron-accepting partners (16, 30, 31), an adaption period of 38-45 days was required for substantial methane to be produced in the initial co-culture. However, with subsequent transfer, ethanol was converted to methane without a substantial lag (Figure 1).

**Figure 1.**
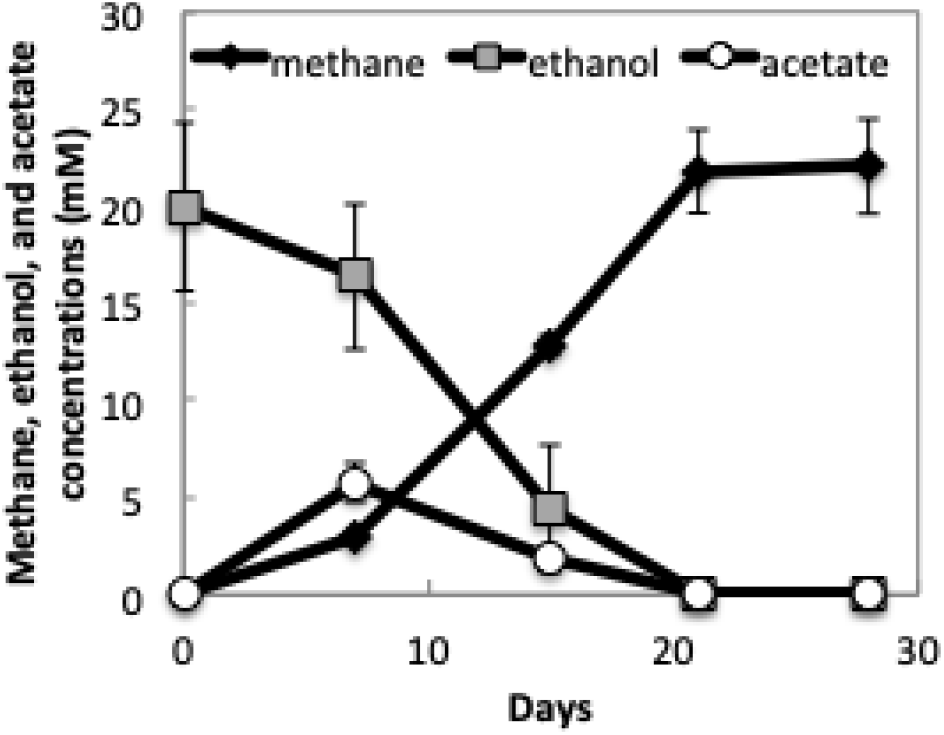
Ethanol consumption and production of methane and acetate in co-cultures established with *G. metallireducens* and *M. acetivorans* after four successive transfers. Data are means and standard deviation of triplicate cultures.

After four transfers of the co-culture, the methane yield was 1.1 mol CH_4_/mol ethanol. Stoichiometric conversion of ethanol to methane yields 1.5 moles of methane, but a portion of the carbon and electrons is required for biomass production. The methane yield in the *G. metallireducens/M. acetivorans* co-culture falls within the range of 0.91 mol CH_4_/mol ethanol to 1.31 mol CH_4_/mol ethanol yields which were obtained when *G. metallireducens* was the electron donating partner for co-cultures grown with other acetotrophic methanogens, such as *Methanothrix harundinacea, M. barkeri, M. mazei, M. vacuolata, M. horonobensis*, and *M. subterranea* (16, 19, 22, 23, 31).

Genes for enzymes specific to the carbon dioxide reduction pathway were more highly expressed in *M. acetivorans* growing in co-culture with *G. metallireducens* versus cells growing in pure culture on acetate (Figure 2, Supplementary Table S1a). This result is in accordance with the fact that carbon dioxide reduction is required to consume the electrons released from ethanol metabolism, accounting for one-third of the methane produced during DIET. Little or no carbon dioxide reduction is expected during growth solely on acetate. H_2_ or formate cannot be the interspecies electron carrier between *G. metallireducens* and *M. acetivorans* for carbon dioxide reduction because *G. metallireducens* cannot grow by metabolizing ethanol with the formation of H_2_ or formate (32) and *M. acetivorans* is unable to use H_2_ or formate as an electron donor (33).

**Figure 2.**
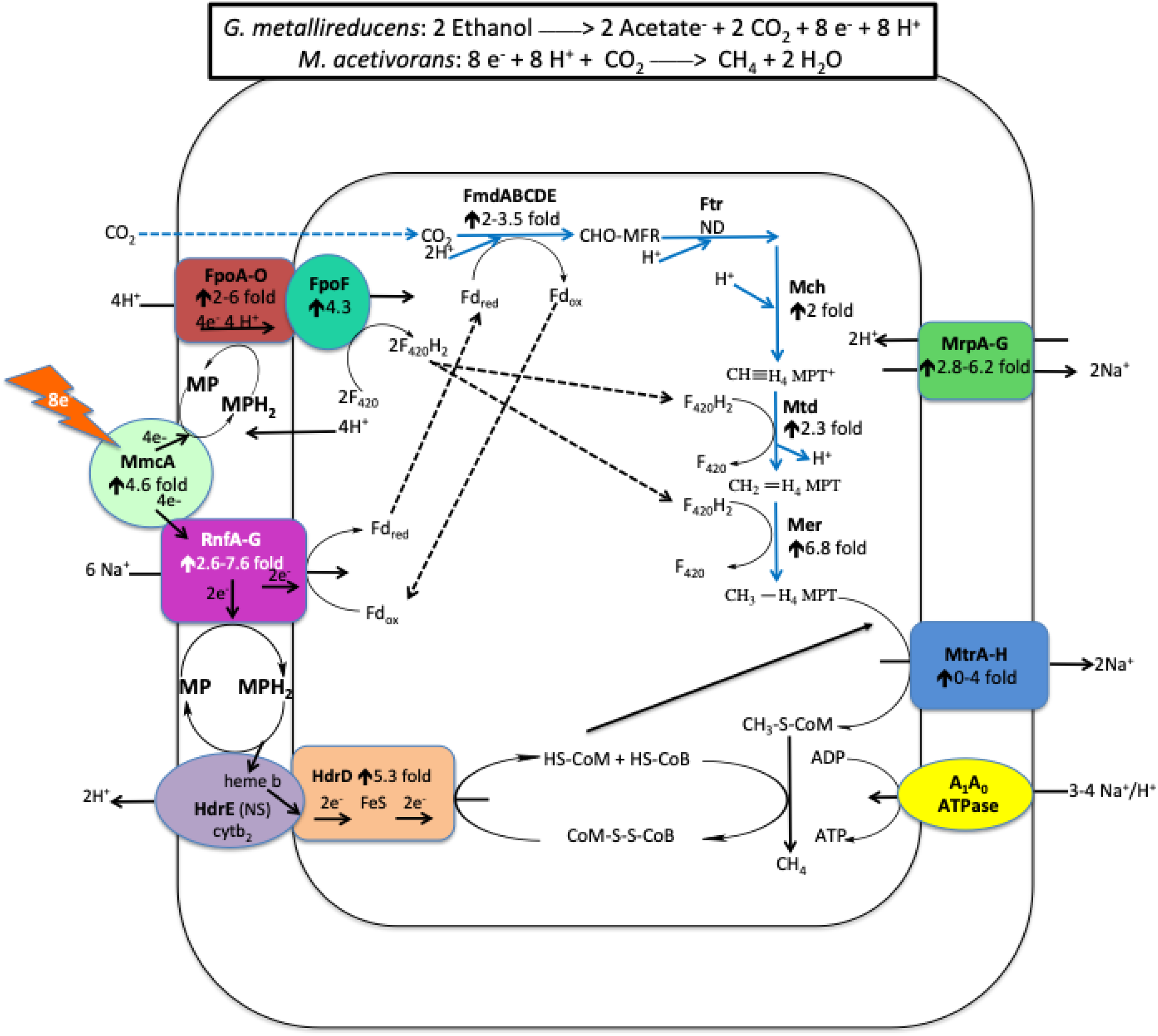
Model for electron and proton flux in *Methanosarcina acetivorans* during direct interspecies electron transfer (DIET) with *Geobacter metallireducens*. Ethanol is provided as the source of electrons and the electron-donating partner (*G. metallireducens*) transfers those electrons to the electron-accepting partner (*M. acetivorans*) for carbon dioxide reduction to CH_4_ through the proposed pathways shown. The degree of increased transcript abundance for subunits of the protein complexes is provided. See main text for more detailed explanation.

Quantitative PCR of DNA extracted from the 4^th^ transfer of triplicate co-cultures with primers targeting the 16S rRNA genes of *G. metallireducens* and *M. acetivorans* revealed that *G. metallireducens* accounted for 60 ± 10% (mean ± standard deviation) of the cells in the co-culture. Confocal and transmission electron microscopy also indicated a near-equal abundance of the two species (Figure 3a,b), and revealed that both species were typically in close proximity, often with more than one cell of each species adjacent to its DIET partner (Figure 3a,b).

**Figure 3.**
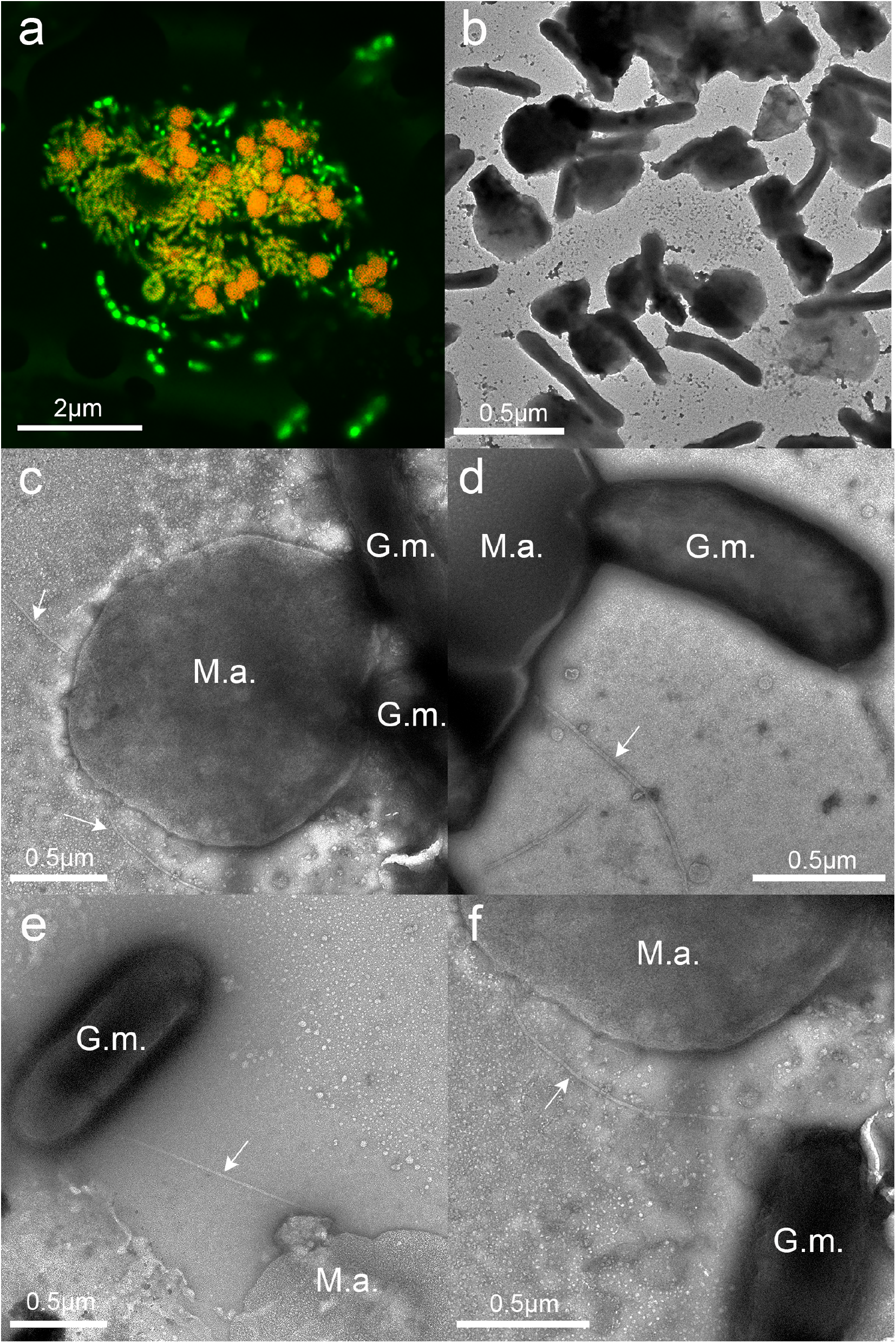
Images of *G. metallireducens/M. acetivorans* co-cultures. (a) Confocal microscopy image demonstrating aggregate size and cell distributions. (b-f) Transmission electron micrographs. Arrows point to archaella extending from the *M. acetivorans* surface. Abbreviations: M.a.: *M. acetivorans*; G.m.: *G. metallireducens*.

### Potential Role(s) for the Archaellum in DIET

Higher magnification TEM images provided further insights into the interactions between *G. metallireducens* and *M. acetivorans* (Figure 3c-e). The outer surfaces of cells of the two species often appeared to be in direct contact (Figure 3c,d). However, there were instances in which filaments (diameter ca. 15 nm), consistent with the appearance of the *M. acetivorans* archaellum (33), appeared to emanate from *M. acetivorans* and connect to juxtaposed cells of *G. metallireducens* (Figure 3e,f).

Genes coding for archaella proteins were not more significantly expressed in DIET-versus acetate-grown cells (Supplementary Table S1) as might be expected because *M. acetivorans* also expresses archaella during growth on acetate (33). In order to evaluate whether the *M. acetivorans* archaella might play a role in DIET, a strain in which two genes for puative archaellin proteins, FlaB1 and FlaB2, were deleted, yielding a strain that did not express archaella (Supplementary Figure S1d). The archaella-deficient strain did not form an effective DIET co-culture with *G. metallireducens* for over 150 days (Figure 4a). However, when the co-cultures were amended with granular activated carbon (GAC), the co-cultures initated with the archaella-deficeint strain produced methane as effectively as co-cultures initated with the parent *M. acetivorans* strain that expressed archaella (Figure 4b).

**Figure 4.**
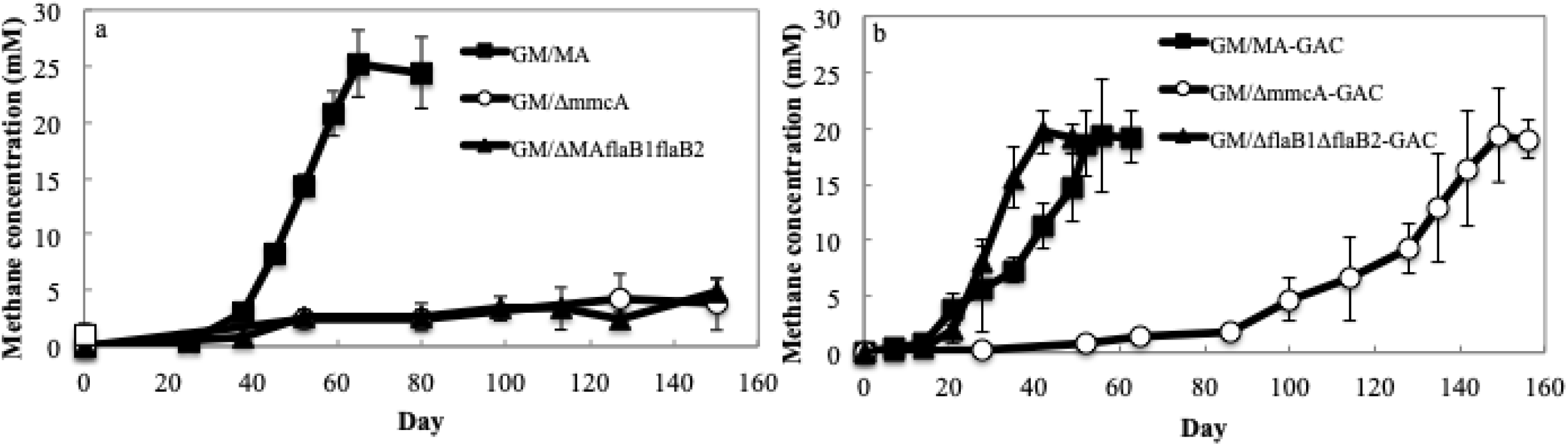
Methane production during initial establishment of co-cultures with *G. metallireducens* and various *M. acetivorans* strains grown with ethanol (20 mM) provided as the electron donor in the absence (a) or presence (b) of granular activated carbon (GAC). Data are means and standard deviation of triplicate cultures. GM: wild-type *G. metallireducens*; MA: wild-type *M. acetivorans*; Δ*mmcA*: *M. acetivorans* strain lacking the gene for the multi-heme cytochrome MmcA; Δ*flaB1*Δ*flaB2*: *M. acetivorans* strain lacking the genes for the archaellins FlaB1 and FlaB2.

GAC and other electrically conductive carbon materials can stimulate DIET between wild-type partners and can enable DIET when genes for key extracellular electron transport proteins that are otherwise essential for DIET, like e-pili, have been deleted (34-37). The DIET partners attach to the GAC rather than each other and the GAC serves as the conduit for long-range interspecies electron transfer (13). Archaella are homologous to type IV pili (38) and the archaellum of *Methanospirillum hungatei* has a conductance 4-fold higher than *G. sulfurreducens* e-pili, demonstrating that at least some archaella can be electrically conductive and might have the potential to be involved in cell-to-cell electron transfer (39). The conductivity of a diversity of e-pili as well as the *Ms. hungatei* archaellum are associated with a high abundance of aromatic amino acids (> 9 %) in the pilin/archaellin monomers and no large gaps (> 40 amino acids) without aromatic amino acids (18). The high density of aromatic amino acids (FlaB1, 11.3%; FlaB2, 9.5%) and the lack of large aromatic-free gaps (largest gaps: FlaB1, 26 amino acids; FlaB2, 29 amino acids) in the *M. acetivorans* archaellins suggest that they might yield conductive archaella. The ability of GAC to rescue the archaella-deficient strain to enable DIET is consistent with a possible archaella role in long-range electron transport. However, other, more traditional roles of archaella, such as conferring motility and facilitating attachment (40) might also help cells locate a DIET partner and/or establish initial interspecies contact. In order to more definitively evaluate a role for the *M. acetivorans* archaellum in interspecies electron transfer it will be necessary to follow the approach employed for evaluating the role of *Geobacter* e-pili in DIET (37) and construct a strain that expresses an archaellum of with potentially low conductivity. However, such studies are technically difficult and well beyond the scope of the current investigation.

### A Role for the Outer-Surface Cytochrome MmcA

Gene expression and deletion studies have indicated that the outer surface multi-heme *c*-type cytochrome MmcA is an important component in *M. acetivorans* for electron transfer to the extracellular electron acceptor AQDS (29). Gene transcripts for MmcA were 5-fold higher (p=0.008) during growth via DIET versus growth on acetate (Figure 2, Supplementary Table S1). Methane production was inhibited in co-cultures initiated with an MmcA-deficient strain of *M. acetivorans* for over 150 days (Figure 4a). These results suggest that MmcA may provide an important route for extracellular electron exchange during DIET.

Unlike the archaella-deficient mutant, GAC did not rapidly rescue the growth of the MmcA-deficient mutant in co-culture (Figure 4b). The co-cultures initiated with the MmcA-deficient strain only grew after a very long lag period. The poor methane production even in the presence of GAC is consistent with the fact that MmcA is thought to be embedded in the membrane of *M. acetivorans* (25, 41). Thus, its role in extracellular electron transfer is expected to be facilitating transmembrane electron transport. Although GAC can enhance long-range electron exchange between the outer cell surface of different species, it does not have a conceivable role in electron transfer across the cell membrane.

*M. acetivorans* has genes for four other putative *c*-type cytochromes, but the presence of these proteins in *M. acetivorans* has yet to be verified and deletion of the genes for each of the four cytochrome genes had no impact on extracellular electron transfer to AQDS even though several of the genes had higher transcript abundance when grown with AQDS as the electron acceptor (29). Transcript abundance for three of these putative cytochrome genes (MA0167, MA2925, MA3739) was higher in DIET-grown *M. acetivorans* than in acetate-grown cells (Supplementary Table S1). Further studies to attempt to document the expression of these proteins in *M. acetivorans* are warranted.

### Potential role for Rnf and Fpo Complexes in DIET

MmcA has the potential to exchange electrons with methanophenazine or the membrane-bound Rnf complex RnfCDGEAB (25, 42-44). Methanophenazine is an important membrane-bound electron carrier and the Rnf complex is physically associated with MmcA in the *M. acetivorans* membrane (25, 41). The Rnf complex oxidizes reduced ferrodoxin with concomitant transport of sodium across the cell membrane from the cell interior to exterior (25, 42). It is proposed that the electrons from ferrodoxin oxidation are transferred directly to methanophenazine during acetotrophic methanogenesis (25, 42, 45) or to MmcA during reduction of extracellular electron acceptors such as Fe(III) and AQDS (25, 29, 46).

Transcripts for the majority of genes coding for Rnf subunits were more significantly expressed in DIET-grown cells than acetate-grown cells (Figure 2, Supplementary Table S2), suggesting an enhanced role for the Rnf complex during DIET. It seems possible that during DIET the Rnf complex functions in the reverse direction proposed for extracellular electron transfer, i.e. accepting electrons to generate the reduced ferrodoxin that is required for the first step in the reduction of carbon dioxide to methane (Figure 2). The most likely electron donor to the Rnf complex is MmcA, which is thought to exchange electrons with Rnf in other forms of *M. acetivorans* electron transfer (25) and, as noted above, is important for DIET (Figure 2). The ferrodoxin reduction requires transfer of sodium to the interior of the cell via the Rnf complex (Figure 2). Ten genes coding for ferredoxin proteins and a gene coding for an unusual flavodoxin (FldA) that can replace ferredoxin as an electron donor under iron-limiting conditions (47) were more than 2 fold more highly expressed (p<0.05) in DIET-grown cells (Supplementary Table S1).

Although the pathway for the biosynthesis of methanophenazine has not been deciphered yet, it resembles respiratory quinones in that it has a polyprenyl side-chain connected to a redox-active moiety (48, 49). Geranylfarnesyl diphosphate is a biosynthetic precursor of methanophenazine, and a homolog (MA0606) of the geranylfarnesyl diphosphate synthase (MM0789) required for methanophenazine biosynthesis in *M. mazei* (50) was 2.42-fold (p=0.01) more highly expressed in DIET grown cells (Supplementary Table S1).

The pathway for carbon dioxide reduction to methane also requires reduced F_420_ (51). The membrane-bound F_420_ dehydrogenase of *M. barkeri* can accept electrons from reduced methanophenazine to generate reduced F_420_ (52) and has been proposed to catalyze F_420_ reduction in a similar manner during *M. barkeri* DIET-based growth (21). This reaction requires concomitant proton translocation from the outside of the cell to the cell interior. Genes for all but one of the Fpo subunits were more highly expressed during *M. acetivorans* growth via DIET versus growth on acetate (Figure 2; Supplementary Table S3). Therefore, electron transfer from MmcA to methanophenazine followed by electron transfer to Fpo is a likely route for generating F_420_H_2_ to support carbon dioxide reduction during DIET (Figure 2).

As in other methanogens, methane production in *M. acetivorans* also requires an electron donor to reduce Coenzyme M 7-mercaptoheptanoylthreonine-phosphate heterodisulfide (CoMS-SCoB) to regenerate coenzyme M (25). It is proposed that during acetoclastic growth the membrane-bound HdrED complex accepts electrons from methanophenazine reduced by the Rnf complex to reduce CoMS-SCoB to HSCoM and HSCoB while pumping two protons from the interior of the cell across the cell membrane (25). Even though the HdrED complex is required for the conversion of acetate to methane, genes for components of this complex were more highly expressed during growth via DIET (Figure 2, Supplementary Table S4). Thus, HdrED is a likely catalyst for CoMS-SCoB reduction (Figure 2). An alternative strategy for reducing CoMS-SCoB is for HdrABC complexes to oxidize F_420_H_2_ in an electron bifurcation reaction that reduces both ferrodoxin and CoMS-SCoB (53, 54). Genes for components of the *M. acetivorans* HdrABC complexes were more highly expressed in DIET-grown cells, suggesting the possibility for multiple routes for electron flux during DIET (Supplementary Table S4).

The proposed route for electron flux during DIET (Figure 2) demonstrates the possibility for energy conservation from carbon dioxide reduction to methane with electrons derived from DIET. The oxidation of two ethanols to acetate and carbon dioxide yields eight electrons required to reduce carbon dioxide to methane. The eight protons that are also generated from this ethanol metabolism must be consumed in order to prevent acidification within the DIET aggregates. Half of these protons are consumed with the proposed Fpo generation of F_420_H_2_ (Figure 2). External sodium ions are needed for the proposed Rnf generation of reduced ferrodoxin. This requirement can be met by the H^+^/Na^+^ antiporter complex (MrpABCDEFG), which adjusts the H^+^/Na^+^ ratio for optimal ATP synthesis by A_1_A_0_ ATP synthase (55, 56). As might be expected, genes for components of this complex are more highly expressed in DIET-grown cells (Figure 2, Supplementary Table S2). The proposed consumption of ten positive charges in the reactions catalyzed by the Fpo and Rnf complexes consumes two more positive charges than the eight that are available from ethanol metabolism. However, the export of two sodiums during the reaction catalyzed by the MtrA-H complex and the two protons exported by HdrED yields a net exterior proton gradient to support ATP generation via ATPase. Detailed functional studies would be required to completely validate this model, but the model is based on previously proposed functions of these *M. acetivorans* components, supporting its feasibility.

### Implications

The results demonstrate that *M. acetivorans* can serve as an electron-accepting partner for DIET and reveal potential outer-surface electrical contacts and routes for electron flux to support DIET-driven carbon dioxide reduction. This is significant because *M. acetivorans*, which is genetically tractable and one of the most intensively studied methanogens (25), is an excellent physiological model for the Type II *Methanosarcina* species that are abundant in many methanogenic soils, sediments, and subsurface environments (22). The results also suggest that different genera of methanogens are likely to employ different strategies for electron uptake during growth via DIET. For example, although MmcA appears to be important for *M. acetivorans* DIET, some *Methanothrix* (31) and *Methanobacterium* (20) species can participate in DIET, but lack *c*-type cytochromes (24).

DIET mechanisms in *M. acetivorans* also appear to differ significantly from those described in *M. barkeri* (21). This is consistent with other substantial differences between Type I (i.e. *M. barkeri*) and Type II (i.e. *M. acetivorans*) *Methanosarcina* species (22). *M. barkeri* lacks MmcA and other *c*-type cytochromes (24). The lack of an Rnf complex in *M. barkeri* requires that electron transport through the membrane to generate reducing equivalents for carbon dioxide reduction relies on the Fpo complex (21).

The diversity of mechanisms for DIET in methanogens suggests that the strategies that rely on gene expression patterns to evaluate the importance of DIET in methanogenic systems will need to accommodate these differences. The mechanisms for extracellular electron exchange in the bacteria and archaea that predominate in anaerobic environments such as soils, sediments, anaerobic digesters, and intestinal systems are still poorly understood (13). For example, although multiple lines of evidence suggest that e-pili are important for extracellular electron transfer in some *Geobacter* species, a model for how e-pili interact with the rest of the *Geobacter* electron transport chain, which could aid in understanding how the archaellum of *M. acetivorans* might ‘plug in’ to membrane electron transport components during DIET, is not yet available (13).

However, the genetic tractability of *M. acetivorans* and the growing information on the biochemistry and function of its key proteins (25); as well as its ability to grow as either an electrogen (transporting electrons to extracellular electron acceptors) (29, 46), or an electrotroph (consuming electrons from an external source), as shown here, suggest that *M. acetivorans* is an excellent model microbe for further study of extracellular electron exchange in archaea.

## Materials and Methods

### Parental strain adaption for co-culture at a compatible salinity

*Geobacter metallireducens* (ATCC 53774) was routinely cultured at 30°C under anaerobic conditions (N_2_:CO_2_, 80:20, vol/vol) with ethanol (20 mM) provided as the electron donor and Fe(III) citrate (56 mM) as the electron acceptor in freshwater medium as previously described (57). *M. acetivorans* strain WWM1 (*Δhpt*) (58), (a gift from William Metcalf at the University of Illinois) was routinely cultured at 37°C in HS-methanol-acetate medium under strict anaerobic conditions as previously described (27, 59).

In order to obtain strains of both microbes that grew at compatible temperatures and salinities, both cultures were adapted to grow at 30°C in MA medium which consisted of the following components per liter: 0.35 g K_2_HPO_4_, 0.23 g KH_2_PO_4_, 0.5 g NH_4_Cl, 4 g NaCl, 1 ml 0.2% wt/vol FeSO_4_, 1 ml trace element solution SL-10 (DSMZ culture collection, medium 320), 10 mM NaHCO_3_, 10 ml Wolin’s vitamin solution (DSMZ culture collection, medium 141), 0.3 mM L-cysteine·HCl, 1 ml 2.7% CaCl_2_·2 H_2_O, and 1 ml 4.5% MgSO_4_·7 H_2_O. The sodium bicarbonate, Wolin’s vitamins, L-cysteine, CaCl_2_, and MgSO_4_ solutions were added from sterile anoxic stocks after the base medium was autoclaved.

For co-culture experiments, *G. metallireducens* and *M. acetivorans* were grown with 20 mM ethanol provided as the electron donor and carbon dioxide as the electron acceptor at 30°C in MA media as previously described (16, 21). For comparative transcriptomic studies *M. acetivorans* was also grown in MA medium with acetate (40 mM) as the sole electron donor.

### *M. acetivorans* mutants

A mutant strain in which the gene for the multi-heme *c*-type cytochrome MmcA was deleted was described previously (29). A strain in which the genes for the archaellin monomer proteins FlaB1 and FlaB2 were deleted was constructed by replacing *flaB1* and *flaB2* genes with *pac* (puromycin resistance gene) (Supplementary Figure S1). The upstream and downstream regions of *flaB1/flaB2* were amplified by PCR with the following primer pairs:

TCTCTCGAGTTCCTTGAAGATATTAAAGGTC/TCTAAGCTTAATGAATCACCTCAATATTGTG and TCTGGATCCAGCTTGAAATCAAACCAC/TCTGCGGCCGCCACTGCAGCTATAA CAC, respectively. The DNA fragments of the upstream and downstream regions were digested with *XhoI/HindIII* and *BamHI/NotI*, respectively. The upstream fragment was ligated with pJK3 (27) and then the downstream fragment was ligated with the pJK3 containing the upstream fragment. The constructed plasmid was linearized with *XhoI* and the linearized plasmid was used for transformation. The deletion of *flaB1*/*flaB2* was verified by PCR with primer pairs, TCTCTCGAGTTCCTTGAAGATATTAAAGGTC (P1)/CCGCCTGCAGTATTCGTTAC (P3) and ACTCTATGCTTGCAGCTGAC (P4)/TCTGCGGCCGCCACTGCAGCTATAACAC (P2) (Supplementary Figure S1). The replacement with the *pac* gene was verified by PCR with a primer pair, AGAGACCCTATCTTACCTGC (P5)/ TCTGCGGCCGCCACTGCAGCTATAACAC (P2) (Supplementary Figure S1). Absence of flagella in the deletion mutant strain was confirmed with transmission electron microscopy (Supplementary Figure S1).

### Analytical techniques

Ethanol in solution was monitored with a gas chromatograph equipped with a headspace sampler and a flame ionization detector (Clarus 600; PerkinElmer Inc., CA). Methane in the headspace was measured by gas chromatography with a flame ionization detector (Shimadzu, GC-8A) as previously described (60). Acetate concentrations were measured with a SHIMADZU high performance liquid chromatograph (HPLC) with an Aminex HPX-87H Ion Exclusion column (300 mm × 7.8 mm) and an eluent of 8.0 mM sulfuric acid.

### Microscopy

Cells were routinely examined by phase-contrast and fluorescence microscopy (BV-2A filter set) with a Nikon E600 microscope. For transmission electron microscopy (TEM), 7 μl of cells were dropcast on plasma-sterilized carbon coated 400 mesh copper ultralight grids for 10 minutes. Liquid was wicked off and the grid was stained with 3 μL 2% uranyl acetate for 15-20 seconds before the liquid was wicked off and air-dried. Transmission electron microscopy was done on a FEI Tecnai 12 at 120kV, spot size 3, with a camera exposure of 200 ms.

Cells for confocal microscopy were harvested (1mL) and vacuumed gently onto a polycarbonate 0.2μm filter, washed with 1mL of wash buffer for 10 minutes, vacuumed, stained with the Live/Dead BacLight Bacterial Viability kit (Thermofisher) (1mL staining, 3 μL each stain per mL) for 10 minutes, vacuumed, destained with wash buffer for 10 minutes, vacuumed, and mounted on glass slides with cover slips and antifade reagent in glycerol. Cells were visualized on a Nikon A1R-SIMe confocal microscope.

### DNA extraction and quantitative PCR

Genomic DNA was extracted from co-cultures with the MasterPure complete DNA purification kit (Lucigen). The proportion of *G. metallireducens* and *M. acetivorans* cells in co-cultures was determined with quantitative PCR using the following primer sets: (i) GS15-16Sq-f (5’-CAGCTCGTGTCGTGAGATGT-3’) and GS15-16Sq-r (5’-GTTTGACACCGGCAGTTTCT-3’) which amplified a 106 bp fragment from the 16S rRNA gene of *G. metallireducens* and (ii) MA-16Sq-f (5’-GTAGTCCCAGCCGTAAACGA-3’) and MA-16Sq-r (5’-CCCGCCAATTCCTTTAAGTT-3’) which amplified a 132 bp fragment of the *M. acetivorans* 16S rRNA gene. Both *G. metallireducens* and *M. acetivorans* have three copies of the 16S rRNA gene in their genomes. Therefore, qPCR results were not influenced by unequal gene copy numbers. Standard curve analysis of both primer sets revealed that they had >95% efficiencies and melt curve analysis yielded a single peak indicating that they were highly specific.

Power SYBR Green PCR Master Mix (Applied Biosystems, Foster City, CA) and an ABI 7500 real-time PCR system were used to amplify and to quantify all PCR products. Each reaction mixture (25 μl) consisted of forward and reverse primers at a final concentration of 200 nM, 5 ng of gDNA, and 12.5 μl of Power SYBR Green PCR Master Mix (Applied Biosystems).

### RNA Extraction

Cells were harvested from triplicate 50 ml cultures of *M. acetivorans* grown alone with acetate (40 mM) provided as a substrate (acetate conditions), or 50 ml cultures of *M. acetivorans* grown in co-culture with *G. metallireducens* with ethanol (20 mM) provided as an electron donor (DIET condition). Cells were harvested during mid-exponential phase when ∼18 mM methane was detected in the headspace.

Cells were split into 50 ml conical tubes (BD Sciences), mixed with RNA Protect (Qiagen) in a 1:1 ratio, and pelleted by centrifugation at 3,000 x g for 15 minutes at 4°C. Pellets were then immediately frozen in liquid nitrogen and stored at -80 °C. Total RNA was extracted from cell pellets as previously described (61), and all six RNA samples (3 acetate, 3 DIET) were cleaned with the RNeasy Mini Kit (Qiagen) and treated with Turbo DNA-free DNase (Ambion). PCR with primers targeting the 16S rRNA gene was then done on all samples to ensure that they were not contaminated with genomic DNA. mRNA was then further enriched from all samples with the MICROB*Express* kit (Ambion), according to the manufacturer’s instructions.

### Illumina sequencing and data analysis

The ScriptSeq™ v2 RNA-Seq Library Preparation Kit (Epicentre) was used to prepare directional multiplex libraries. Paired end sequencing was then performed on these libraries with a Hi-Seq 2000 platform at the Deep Sequencing Core Facility at the University of Massachusetts Medical School in Worchester, Massachusetts.

Raw data was quality checked with FASTQC (http://www.bioinformatics.babraham.ac.uk/projects/fastqc/), and initial raw non-filtered forward and reverse sequencing libraries contained an average of 68,911,030 +/- 21,863,730 reads that were ∼100 basepairs long (Supplementary Table S5). Sequences from all of the libraries were trimmed and filtered with Trimmomatic (62) which yielded an average of 55,239,290 +/- 29,060,121 quality reads per RNAseq library.

All paired-end reads were then merged with FLASH (63), resulting in 32,159,242 +/- 22,219,390 reads with an average read length of 134 +/- 28 basepairs. Ribosomal RNA (rRNA) reads were then removed from the libraries with SortMeRNA (64), which resulted in 4,959,312 +/- 2,340,361 mRNA reads.

### Mapping of mRNA reads

Trimmed and filtered mRNA reads from the triplicate samples for the two different culture conditions were mapped against the *M. acetivorans* strain C2A genome (NC_003552) downloaded from IMG/MER (img.jgi.doe.gov) using ArrayStar software (DNAStar). Reads were normalized and processed for differential expression studies using the edgeR package in Bioconductor (65). All genes that were ≥2 fold differentially expressed with p-values ≤ 0.05 are reported in Supplementary Table S1.

## Data Availability

Illumina sequence reads have been submitted to the SRA NCBI database under BioProject PRJNA727272 and Biosamples SAMN19011637, and SAMN19011638.

## Acknowledgments

The confocal microscopy data was gathered in the Light Microscopy Facility and Nikon Center of Excellence at the Institute for Applied Life Sciences, UMass Amherst with support from the Massachusetts Life Sciences Center. The transmission electron microscopy images were collected in the Electron Microscopy Facility at the Institute for Applied Life Sciences, UMass Amherst.

This research was supported by the Army Research Office and was accomplished under Grant Number W911NF-17-1-0345. The views and conclusions contained in this document are those of the authors and should not be interpreted as representing the official policies, either expressed or implied, of the Army Research Office or the U.S. Government.

## Ethics declarations

The authors do not declare any conflicts of interest.

## Notes

### Competing Interest Statement

The authors have declared no competing interest.

