## Supplemental Figure S1 for "Mechanisms for Electron Uptake by *Methanosarcina acetivorans* During Direct Interspecies Electron Transfer"

### Slide 1
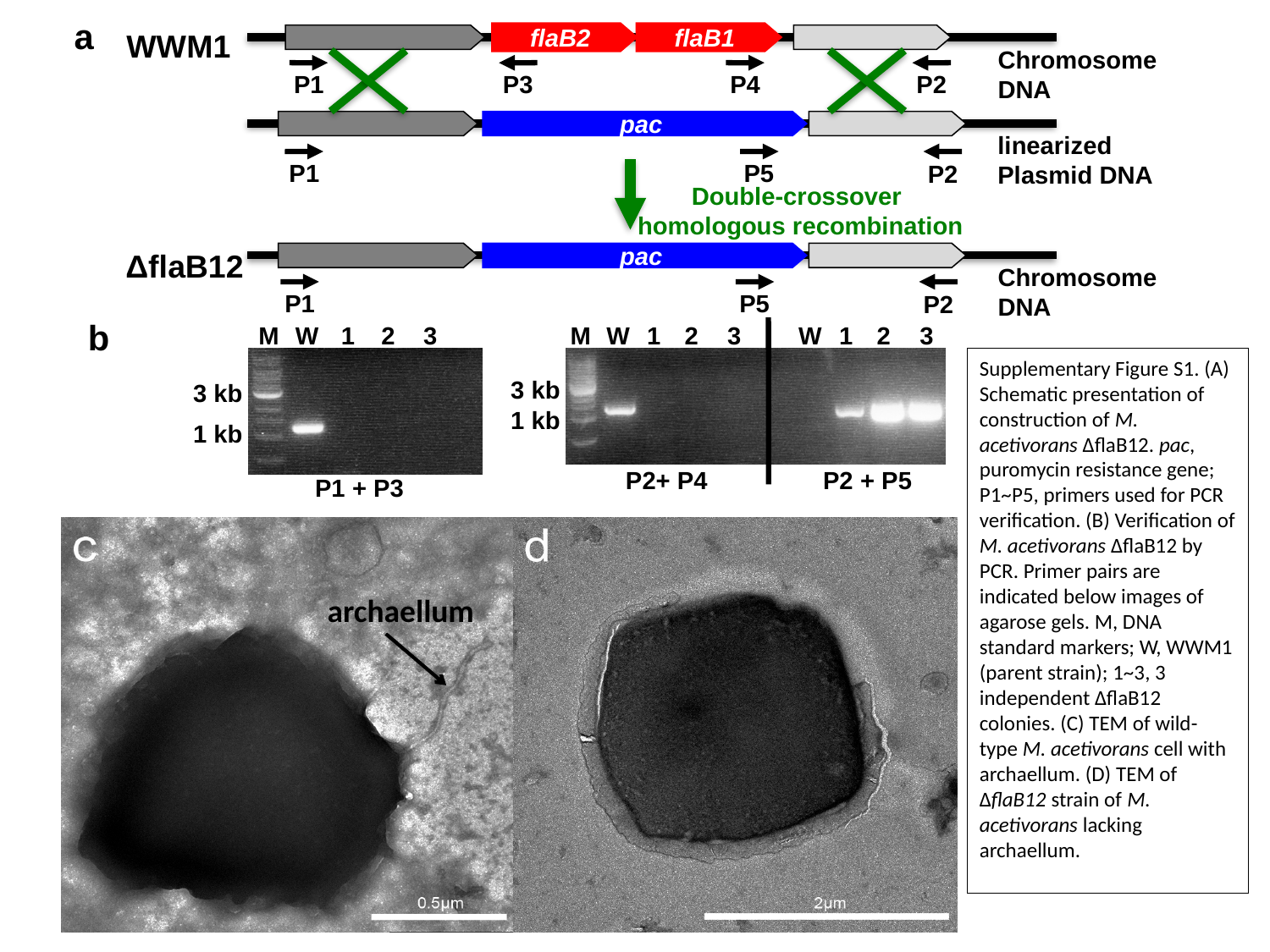

a
WWM1
flaB2
flaB1
Chromosome
DNA
P4
P3
P1
P2
pac
linearized
Plasmid DNA
P5
P1
P2
Double-crossover
homologous recombination
ΔflaB12
pac
Chromosome
DNA
P5
P1
P2
b
M
W
1
2
3
M
W
1
2
3
W
1
2
3
3 kb
3 kb
1 kb
1 kb
P2+ P4
P2 + P5
P1 + P3
Supplementary Figure S1. (A) Schematic presentation of construction of M. acetivorans ∆flaB12. pac, puromycin resistance gene; P1~P5, primers used for PCR verification. (B) Verification of M. acetivorans ∆flaB12 by PCR. Primer pairs are indicated below images of agarose gels. M, DNA standard markers; W, WWM1 (parent strain); 1~3, 3 independent ∆flaB12 colonies. (C) TEM of wild-type M. acetivorans cell with archaellum. (D) TEM of ∆flaB12 strain of M. acetivorans lacking archaellum.
archaellum
