## Supplemental Table S2 for "Mechanisms for Electron Uptake by *Methanosarcina acetivorans* During Direct Interspecies Electron Transfer"

Supplementary Table S2. Genes coding for subunits from Rnf and Mrp complexes that were at least 2-fold more significantly expressed in DIET grown *M. acetivorans* cells (grown in co-culture with *G. metallireducens*) compared to *M. acetivorans* cells grown with acetate as the sole substrate for methanogenesis.

Genes were only considered differentially expressed if the p-values were ≤0.05.

NS: no significant difference in read abundance

| Locus ID | Annotation | Gene | Fold up-regulated DIET vs acetate | p-value |
| --- | --- | --- | --- | --- |
| MA0659 | electron transport complex protein RnfC | *rnfC* | 2.55 | 0.03 |
| MA0660 | electron transport complex protein RnfD | *rnfD* | NS | - |
| MA0661 | electron transport complex protein RnfG | *rnfG* | 5.01 | 0.001 |
| MA0662 | electron transport complex protein RnfE | *rnfE* | 3.26 | 0.006 |
| MA0663 | electron transport complex protein RnfA | *rnfA* | 7.55 | 0.0009 |
| MA0664 | electron transport complex protein RnfB | *rnfB* | 3.29 | 0.01 |
| MA1799 | FldA Flavodoxin | *fldA* | 2.48 | 0.03 |
| MA4572 | multisubunit sodium/proton antiporter, MrpA subunit | *mrpA* | 6.15 | 0.006 |
| MA4665 | multisubunit sodium/proton antiporter, MrpB subunit | *mrpB* | NS | - |
| MA4570 | multisubunit sodium/proton antiporter, MrpC subunit | *mrpC* | 5.33 | 0.001 |
| MA4569 | multisubunit sodium/proton antiporter, MrpD subunit | *mrpD* | 2.76 | 0.04 |
| MA4568 | multisubunit sodium/proton antiporter, MrpE subunit | *mrpE* | NS | - |
| MA4567 | multisubunit sodium/proton antiporter, MrpF subunit | *mrpF* | 5.91 | 0.004 |
| MA4566 | multisubunit sodium/proton antiporter, MrpG subunit | *mrpG* | 3.19 | 0.02 |
