## Supplemental Table S3 for "Mechanisms for Electron Uptake by *Methanosarcina acetivorans* During Direct Interspecies Electron Transfer"

Supplementary Table S3. Genes coding for subunits from the Fpo dehydrogenase complex that were at least 2-fold more significantly expressed in DIET grown *M. acetivorans* cells (grown in co-culture with *G. metallireducens*) compared to *M. acetivorans* cells grown with acetate as the sole substrate for methanogenesis.

Genes were only considered differentially expressed if the p-values were ≤0.05.

NS: no significant difference in read abundance

| Locus ID | Annotation | Gene | DIET vs acetate | p-value |
| --- | --- | --- | --- | --- |
| MA1495 | F_420_H_2_ dehydrogenase subunit A | *fpoA* | NS | - |
| MA1496 | F_420_H_2_ dehydrogenase subunit B | *fpoB* | 4.48 | 0.0006 |
| MA1497 | F_420_H_2_ dehydrogenase subunit C | *fpoC* | 2.91 | 0.003 |
| MA1498 | F_420_H_2_ dehydrogenase subunit D | *fpoD* | 3.36 | 0.002 |
| MA1499 | F_420_H_2_ dehydrogenase subunit H | *fpoH* | 5.98 | 0.0002 |
| MA1500 | F_420_H_2_ dehydrogenase subunit I | *fpoI* | 3.68 | 0.004 |
| MA1501 | F_420_H_2_ dehydrogenase subunit J | *fpoJ1* | 4.82 | 0.0003 |
| MA1502 | F_420_H_2_ dehydrogenase subunit J | *fpoJ2* | 5.69 | 0.0007 |
| MA1503 | F_420_H_2_ dehydrogenase subunit K | *fpoK* | 2.03 | 0.01 |
| MA1504 | F_420_H_2_ dehydrogenase subunit L | *fpoL* | 3.21 | 0.002 |
| MA1505 | F_420_H_2_ dehydrogenase subunit M | *fpoM* | 4.44 | 0.001 |
| MA1506 | F_420_H_2_ dehydrogenase subunit N | *fpoN* | 6.57 | 0.001 |
| MA1507 | F_420_H_2_ dehydrogenase subunit 0 | *fpoO* | 3.26 | 0.005 |
| MA3732 | F_420_H_2_ dehydrogenase subunit F | *fpoF* | 4.31 | 0.0008 |
