## Supplemental Table S4 for "Mechanisms for Electron Uptake by *Methanosarcina acetivorans* During Direct Interspecies Electron Transfer"

**Supplementary Table S4**. Differential expression of genes coding for various subunits from the membrane bound heterodisulfide reductase complex (HdrDE) and the two soluble heterodisulfide reductase complexes (HdrABC). Genes were only considered differentially expressed if there was at least a two-fold difference and p values were ≤ 0.05.

| Locus ID | Gene annotation | Gene name | Fold up-regulated in DIET vs acetate | p-value |
| --- | --- | --- | --- | --- |
| MA0687 | Heterodisulfide reductase subunit E | *hdrE* | NS | NS |
| MA0688 | Heterodisulfide reductase subunit D | *hdrD* | 5.28 | 0.003 |
| MA2868 | CoB--CoM heterodisulfide reductase subunit A | *hdrA* | 2.54 | 0.009 |
| MA3128 | CoB--CoM heterodisulfide reductase subunit A | *hdrA* | 7.13 | 0.03 |
| MA4237 | CoB--CoM heterodisulfide reductase subunit B | *hdrB* | 2.28 | 0.02 |
| MA4236 | CoB--CoM heterodisulfide reductase subunit C | *hdrC* | NS | NS |
| MA3126 | heterodisulfide reductase, subunit B | *hdrB* | 4.02 | 0.05 |
| MA3127 | heterodisulfide reductase, subunit C | *hdrC* | NS | NS |
| MA0526 | heterodisulfide reductase, subunit D | *hdrD* | 2.53 | 0.007 |
