## Supplemental Table S5 for "Mechanisms for Electron Uptake by *Methanosarcina acetivorans* During Direct Interspecies Electron Transfer"

Supplementary Table S5A. Summary of statistics from RNAseq libraries assembled from RNA extracted from *Methanosarcina* *acetivorans* cells grown via acetotrophic methanogenesis with 40 mM acetate provided as the substrate (acetate condition; 3 biological replicates).

|  | Mean | SEM | Max | Min | Units |
| --- | --- | --- | --- | --- | --- |
| Raw (unfiltered) reads | 53,451,038 | 4,661,888 | 63,336,930 | 39,037,040 | #reads/library |
| QC-filtered reads | 34,690,681 | 3,833,156 | 42,104,990 | 24,133,014 | #reads/library |
| QC-filtered merged paired end reads | 16,447,761 | 2,954,066 | 11,110,961 | 21,311,254 | #reads/library |
| QC-filtered merged paired mRNA reads | 3,304,427 | 564,875 | 4,081,120 | 2,205,580 | #mRNA reads/library |
| QC-filtered merged paired end read size | 114.5 | 8.19 | 190 | 99 | Bases/read |

Supplementary Table S5B. Summary of statistics from RNAseq libraries assembled from RNA extracted from co-cultures of *Methanosarcina* *acetivorans* and *Geobacter metallireducens* cells grown with ethanol (20 mM) provided as the electron donor for DIET (DIET condition; 3 biological replicates).

|  | Mean | SEM | Max | Min | Units |
| --- | --- | --- | --- | --- | --- |
| Raw (unfiltered) reads | 84,371,021 | 2,983,587 | 92,482,194 | 76,141,695 | #reads/library |
| QC-filtered reads | 75,787,898 | 1,531,665 | 80,600,666 | 72,909,349 | #reads/library |
| QC-filtered merged paired end reads | 47,870,723 | 2,656,797 | 51,201,184 | 42,619,863 | #reads/library |
| QC-filtered merged paired mRNA reads | 6,614,197 | 587,282 | 7,728,580 | 5,735,580 | #mRNA reads/library |
| QC-filtered merged paired end read size | 154.37 | 1.05 | 190 | 100 | Bases/read |
